# Tissue-specific metabolomic signatures for a *doublesex* model of reduced sexual dimorphism

**DOI:** 10.1101/2024.09.11.612537

**Authors:** Rene Coig, Benjamin R. Harrison, Richard S. Johnson, Michael J. MacCoss, Daniel E.L. Promislow

**Affiliations:** Department of Laboratory Medicine and Pathology, University of Washington, Seattle, Washington, United States; Department of Genome Sciences, University of Washington, Seattle, WA, United States; Department of Biology, University of Washington, Seattle, WA, United States; Jean Mayer USDA Human Nutrition Research Center on Aging, Tufts University, Boston, MA, United States

**Keywords:** *Drosophila melanogaster*, sexual dimorphism, sex differences, metabolism

## Abstract

Sex has a major effect on the metabolome. However, we do not yet understand the degree to which these quantitative sex differences in metabolism are associated with anatomical dimorphism and modulated by sex-specific tissues. In the fruit fly, *Drosophila melanogaster*, knocking out the *doublesex* (*dsx*) gene gives rise to adults with intermediate sex characteristics. Here we sought to determine the degree to which this key node in sexual development leads to sex differences in the fly metabolome. We measured 91 metabolites across head, thorax and abdomen in *Drosophila*, comparing the differences between distinctly sex-dimorphic flies with those of reduced sexual dimorphism: *dsx* null flies. Notably, in the reduced dimorphism flies, we observed a sex difference in only 1 of 91 metabolites, kynurenate, whereas 51% of metabolites (46/91) were significantly different between wildtype XX and XY flies in at least one tissue, suggesting that *dsx* plays a major role in sex differences in fly metabolism. Kynurenate was consistently higher in XX flies in both the presence and absence of functioning *dsx*. We observed tissue-specific consequences of knocking out *dsx*. Metabolites affected by sex were significantly enriched in branched chain amino acid metabolism and the mTOR pathway. This highlights the importance of considering variation in genes that cause anatomical sexual dimorphism when analyzing sex differences in metabolic profiles and interpreting their biological significance.

## Introduction

Sex differences in biological processes are pervasive and have far-reaching implications for health and disease (1–3). In *Drosophila*, these differences extend beyond reproductive functions and include many aspects of cellular metabolism (4–8). In sex differences research, a distinction can be made between sexual dimorphism (the categorical, anatomical difference in morphology between the sexes) and sex differences (the quantitative, bimodal distribution of quantitative traits between the sexes) (9). Currently, there is a significant gap in our understanding of how variation in anatomical sexual dimorphism impacts quantitative sex differences in metabolism.

The metabolome is the entire spectrum of small-molecule metabolites found within a biological sample (10). These metabolites, such as lipids, amino acids, nucleotides, and other small molecules, offer a snapshot of an organism’s physiological state (11,12). Metabolome profiles can vary significantly between sexes (13–15) and genetic factors influence the *Drosophila* metabolome (16–19). Thus, genes that lead to sexually dimorphic anatomy might also lead to sex differences in metabolite levels. Further, variants in sex development genes that increase or decrease the degree of anatomical dimorphism may lead to similar increases or decreases in the effects of sex observed in the metabolome.

Different tissues in an organism perform specialized functions, and their metabolic needs and activities are tailored to support these functions. For example, muscle tissue requires energy for contraction and movement, while liver tissue is involved in detoxification and glucose metabolism. Sex-specific tissues such as ovaries or testes may have unique metabolic demands to perform their specialized functions related to reproduction. Further, communication between reproductive organs and other tissues may influence the overall metabolic landscape of the body. For example, 20- hydroxyecdysone circulating in the hemolymph can influence *Drosophila* oocyte development (20). Currently, there is a gap in our understanding of how the presence, absence or variation in the function of sex-specific tissues such as ovaries or testes may affect observed sex differences in the fly metabolome.

This study aims to address these gaps by measuring the tissue-specific metabolome in a *Drosophila* model of reduced sexual dimorphism. The *Drosophila* sex development pathway is well characterized (21,22), providing an excellent system for genetic manipulation of sexual dimorphism. In *Drosophila*, the *doublesex* (*dsx*) gene acts in a global alternative splicing cascade, which results in female- and male-specific protein isoforms of *dsx* (DSX^F^ and DSX^M^) that are essential for determining the sexual fate of cells throughout the organism (23–25). Knockout of *dsx* disrupts this cascade, leading to incomplete development of gonads and genitals (26). *Dsx* is also known to function as a tissue-specific regulator throughout development (27–29), not just in gonads and genitals but in seemingly non-dimorphic tissues such as legs (30). Throughout adulthood, *dsx* gene products continue to be expressed in the gonads (31), as well as the head (32), particularly in the central nervous system (33,34), and in the fat body (27). This makes *dsx* an exceptionally powerful gene for unraveling the tissue-specific metabolic consequences of sexual dimorphism in *Drosophila*.

In this study, we aim to compare metabolome profiles of distinctly sex-dimorphic wildtype flies with those of reduced sexual dimorphism *dsx* null flies across head, thorax and abdominal tissues using LC-MS targeted metabolomic analysis. We hypothesized that reduced dimorphism in *dsx* null flies would be associated with a reduction in sex differences observed in the metabolome. The primary focus of this analysis is to evaluate how sex differences in the metabolome of wildtype flies, as compared to sex differences in the metabolome of *dsx* null flies, vary. We thus refer to each group of XX and XY flies (wildtype or *dsx* null) as a sex difference group (SD group) and compare the effect sizes of sex differences in the metabolome between these SD groups. Our findings reveal that sex differences in the metabolome are greatly reduced between *dsx* null sexes as compared to wildtype sexes. While *dsx* knockout led to a loss of significant sex differences in almost all metabolite features, the specific features that were significantly different between wildtype sexes varied among the three tissues. These results highlight the importance of considering anatomical sexual dimorphism in metabolomic studies, contributing to a better understanding of sex differences in biological processes.

## Results

We used LC-MS to measure metabolite concentrations in head, thorax and abdomen tissues of four types of flies in a *w*^1118^ genetic background: XX-wildtype, XX-*dsx* null, XY- *dsx* null and XY-wildtype flies. We first report and compare sex differences observed in the global metabolome for both SD groups. Next, we report the tissue-specificity of these sex differences. Last, we report pathway analysis results for enrichment among metabolites with significant effects of sex.

### Reduction in sexual dimorphism is accompanied by a reduction in global sex differences in the metabolome

The primary purpose of this study was to query the degree to which anatomical dimorphism might impact sex differences in metabolism. We hypothesized that global sex differences in the metabolome of *dsx* null flies would be significantly reduced from that of wildtype flies. Principal component analysis (PCA) of all samples together strongly separated tissues from one another, with PC1 segregating abdomen samples from both head and thorax, and PC2 discriminating head from thorax (Supplementary Fig 1). We thus analyzed the data from each tissue separately.

Within each tissue, PCA separated all four genotypes from each other, with the *dsx* null metabolome profiles largely intermediate to those of wildtype flies on PC1 in the thorax and abdomen and in PC2 for head tissue (Fig 1, Supplementary Table 1A). However, on other PCs, the metabolomes of *dsx* null flies were outside the ranges of wildtype sexes (such as PC1 in head and PC2 in thorax), indicating that no single ruler can measure sex differences in the metabolome directly from “female” to “male” in *Drosophila*. Notably, there was little overlap in the metabolomes of all four groups of flies regardless of tissue.

**Fig 1.**
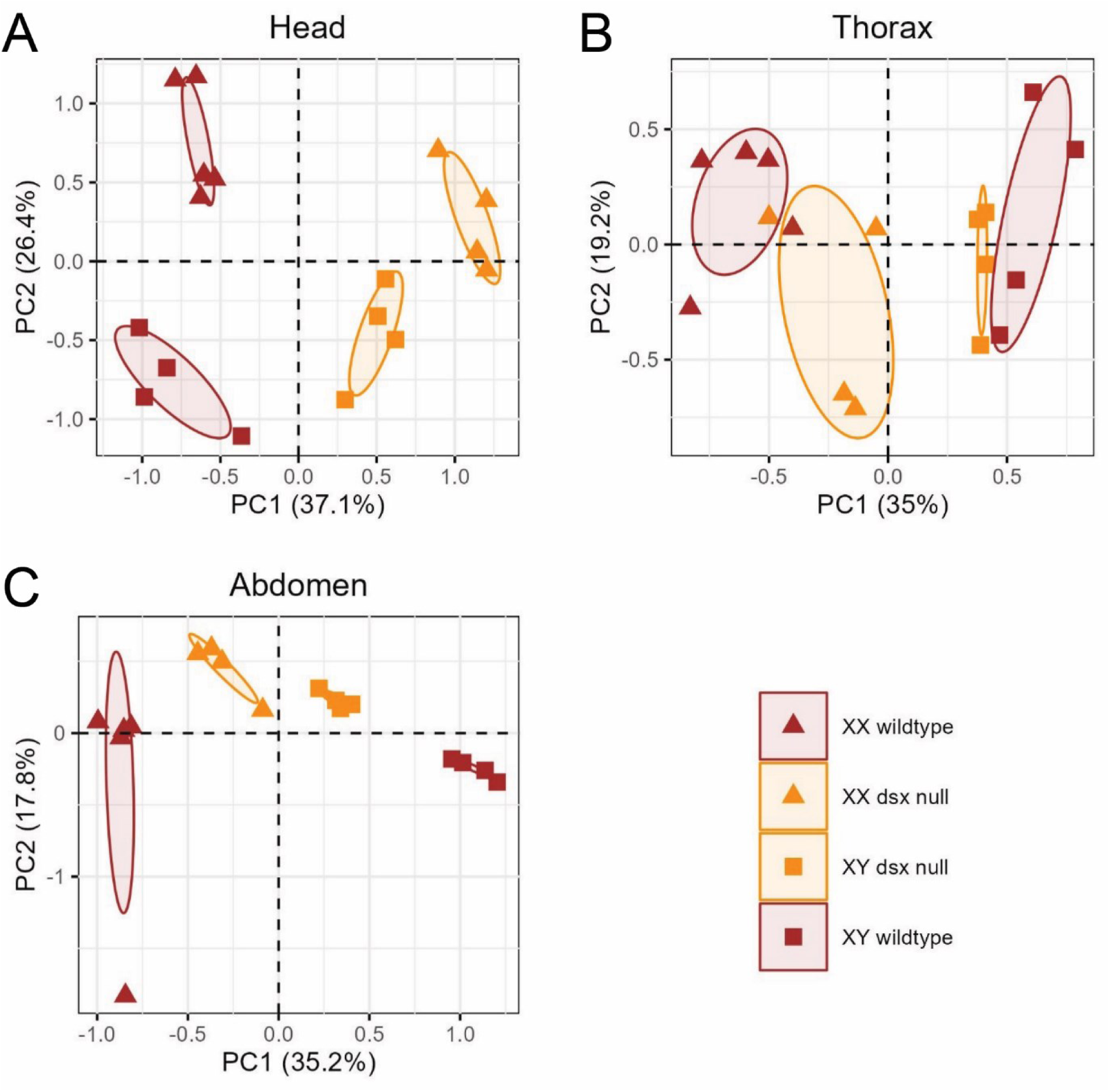
PCA of metabolome samples by tissue. PCA for (A) head tissue, (B) thorax tissue and (C) abdomen tissue. *Dsx* null flies are colored in orange, wildtype flies are colored in brown. Triangles represent XX and squares represent XY fly samples.

We next analyzed sex differences between SD groups independently for each metabolite using ANOVA. As expected, we observed strong sex differences in the metabolome of wildtype flies, with 46 metabolites (51%) at FDR<5% in at least one tissue (Table 1, Supplementary Table 1B). We refer to these metabolites as “SD metabolites”. We found no indication of a sex bias in the directionality of SD metabolites with higher levels in XX as compared to XY (Table 2). Comparisons of the magnitude of sex effect sizes among the SD metabolites in SD groups confirmed that sex differences were significantly reduced in the *dsx* null head (p = 0.002), thorax (p = 3E-04) and abdomen (p = 1E-06) (Fig 2A). Including all metabolites in the analysis, the reduction was less significant; head (p = 0.007), thorax (p = 0.02), abdomen (p = 6E-04), (Fig 2B).

**Fig 2.**
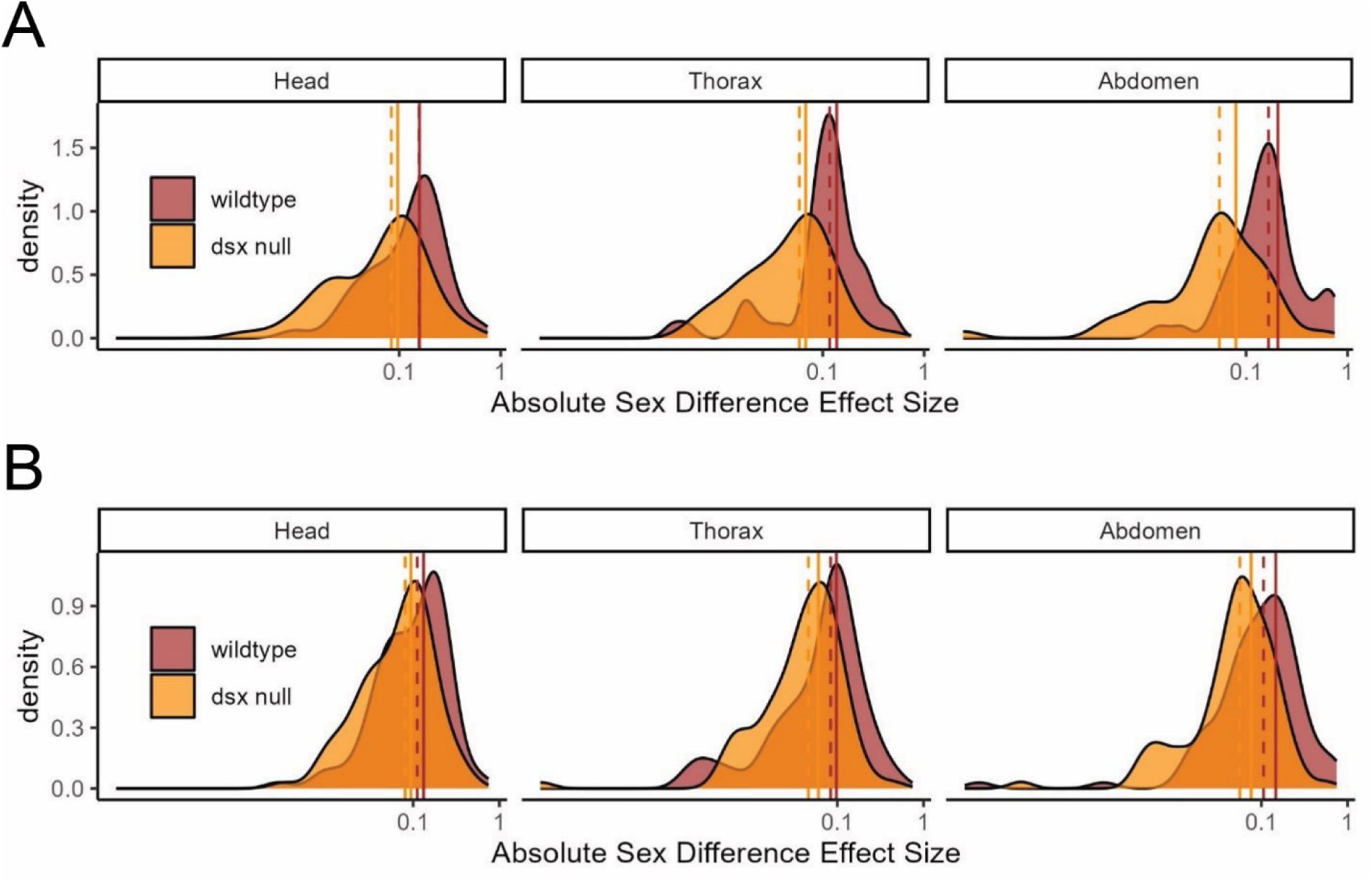
Density of sex effect sizes across metabolites in wildtype and *dsx* null sexes. (A) Density plot across SD metabolites, those metabolites with a significant effect of sex in wildtype flies (N = 46 metabolites). Sex effect sizes across metabolites are significantly different between wildtype and *dsx* null groups: head (p = 0.002), thorax (p = 3E-04) and abdomen (p = 1E-06). (B) Density plot across all metabolites (N = 91 metabolites). Sex effect sizes across metabolites are significantly different between wildtype and *dsx* null groups: head (p = 0.007), thorax (p = 0.02), abdomen (p = 6E-04). Vertical lines mark the median (dashed) and mean (solid) effect size across metabolites. Effect sizes are plotted as absolute values on a log10 scale.

**Table 1.** Summary of sex effect magnitude across metabolites by tissue.

|  |  | **N**<br>**Metabolites**<br>**FDR <5%** | **%**<br>**Metabolites**<br>**FDR <5%** | **Mean ES**<br>**Metabolites**<br>**FDR <5%** | **Mean ES**<br>**All**<br>**Metabolites** |
| --- | --- | --- | --- | --- | --- |
| wildtype | Head | 22 | 24% | 0.16 | 0.13 |
|  | Thorax | 33 | 36% | 0.14 | 0.10 |
|  | Abdomen | 24 | 26% | 0.21 | 0.15 |
| *dsx* null | Head | 1 | 1% | 0.10 | 0.09 |
|  | Thorax | 1 | 1% | 0.07 | 0.06 |
|  | Abdomen | 1 | 1% | 0.08 | 0.08 |

Metabolites with a significant effect between sexes (referred to as SD metabolites) at FDR <5%. Effect size (ES) is calculated using the TukeyHSD function in R.

**Table 2.** Summary of sex effect directionality across tissues.

|  | **N Metabolites**<br>**Higher in XX flies** | **N Metabolites**<br>**Higher in XY flies** |
| --- | --- | --- |
| Head | 10 | 12 |
| Thorax | 15 | 18 |
| Abdomen | 11 | 13 |

Number of metabolites with higher levels in XX as compared to XY wildtype flies.

Of all SD metabolites, only kynurenate maintained a significant sex difference in both wildtype and *dsx* null flies with elevated levels in XX flies of both SD groups across all three tissues (Table 3). No other metabolites were significantly different between XX and XY flies in the *dsx* null SD group.

**Table 3.** Tissue-consistent SD metabolites.

| Head | Effect Size |  | Adjusted p-value (FDR) |  |
| --- | --- | --- | --- | --- |
|  | wildtype | *dsx* null | wildtype | *dsx* null |
| 1-METHYL-L-HISTIDINE | 0.16 | 0.07 | 0.0003 | 0.3 |
| DEOXYCARNITINE | -0.23 | 0.07 | 3.1E-05 | 0.4 |
| KYNURENATE | -0.52 | -0.46 | 4.7E-09 | 5.7E-08 |
| N-METHYLGLUTAMATE | 0.19 | 0.15 | 0.01 | 0.2 |
| ADMA | -0.22 | -0.03 | 0.002 | 1 |
| TYROSINE | -0.23 | -0.01 | 0.05 | 1 |
| VALINE | -0.24 | -0.09 | 1.4E-05 | 0.2 |
| Thorax | Effect Size |  | Adjusted p-value (FDR) |  |
|  | wildtype | *dsx* null | wildtype | *dsx* null |
| 1-METHYL-L-HISTIDINE | 0.15 | 0.03 | 9.5E-06 | 0.7 |
| DEOXYCARNITINE | -0.17 | 0.09 | 0.0001 | 0.2 |
| KYNURENATE | -0.44 | -0.44 | 5.3E-06 | 2.3E-05 |
| N-METHYLGLUTAMATE | 0.13 | 0.07 | 0.004 | 0.4 |
| ADMA | -0.26 | 0.01 | 0.0003 | 1 |
| TYROSINE | -0.12 | -0.03 | 0.0002 | 0.9 |
| VALINE | -0.24 | -0.11 | 1.9E-05 | 0.1 |
| Abdomen | Effect Size |  | Adjusted p-value (FDR) |  |
|  | wildtype | *dsx* null | wildtype | *dsx* null |
| 1-METHYL-L-HISTIDINE | 0.08 | 0.05 | 0.03 | 0.6 |
| DEOXYCARNITINE | -0.17 | 0.13 | 0.01 | 0.2 |
| KYNURENATE | -0.68 | -0.56 | 2.1E-06 | 4.1E-05 |
| N-METHYLGLUTAMATE | 0.14 | 0.08 | 0.04 | 0.8 |
| ADMA | -0.34 | -0.06 | 0.0003 | 1 |
| TYROSINE | -0.14 | -0.05 | 0.004 | 0.9 |
| VALINE | -0.17 | -0.05 | 0.004 | 1.0 |

Effect size is calculated between XX and XY flies using TukeyHSD function in R. A positive effect size indicates values are higher in XY flies. ADMA refers to N,N-dimethyl-arginine.

### *Dsx* influences sex differences in the metabolome in a tissue-specific manner

Across tissues, the number and identity of SD metabolites varied (head = 22, thorax = 33, abdomen = 24), with thorax tissue sharing more SD metabolites with both the head and abdomen than the head and abdomen shared with each other (Fig 3A). Each tissue also had unique SD metabolites, which were present but did not show sex differences at FDR<5% in the other tissues. SD metabolites unique to the head included 3-nitro-L-tyrosine, gamma-aminobutyrate, and nicotinamide mononucleotide. The thorax tissue had eight unique SD metabolites: agmatine sulfate, amino-isobutyrate, deoxy- guanosine, guanosine, histamine, pantothenate, phosphorylcholine (aka phosphocholine), and putrescine. Abdomen tissue had nine unique SD metabolites, including 4-imidazoleacetate, hippurate, hypoxanthine, kynurenine, L-carnitine, O-acetylcarnitine, ophthalmate, proline, and spermidine. Seven metabolites were significantly different between wildtype sexes across all tissues (Fig 3B). These features are 1-methyl-L-histidine, deoxycarnitine, kynurenate, N-methyl-glutamate, N,N-dimethyl- arginine, tyrosine and valine.

**Fig 3.**
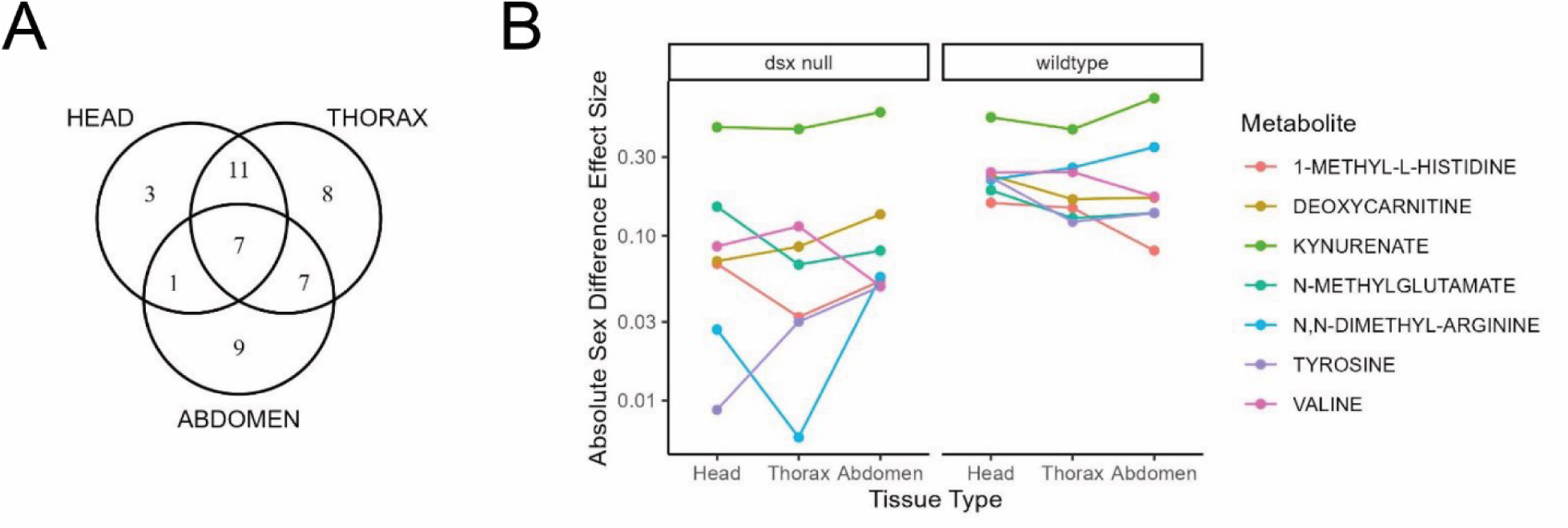
**Tissue specificity of SD metabolites**. (A) Venn diagram of number of SD metabolites that overlap across tissues. (B) Sex effect sizes for seven SD metabolites significantly different between wildtype sexes across all three tissues. Effect sizes are calculated from the TukeyHSD function in R. Kynurenate, in light green, is the one metabolite that maintains a significant sex difference in both wildtype and *dsx* null flies across all three tissues. Effect sizes are plotted on a log10 scale.

Overall, we found that the influence of *dsx* on sex differences in the metabolome is highly tissue specific. Some metabolites showed sex differences consistently across all tissues while sex differences in other metabolites were unique to specific tissues. This tissue specificity could provide insights into how metabolic sex differences contribute to the diverse physiological and behavioral traits observed between sexes.

### SD metabolites are enriched in cellular growth and energy metabolism pathways

We next conducted pathway enrichment analysis for all SD metabolites and identified 11 pathways with significant regulatory differences between wildtype sexes at FDR<5% (Supplementary Table 1C). The top 5 most significantly enriched pathways included phototransduction (KEGG ID: dme04745), branched-chain amino acid (BCAA) degradation (KEGG ID: dme00280), fatty acid elongation (KEGG ID: dme00062), dorso- ventral axis formation (KEGG ID: dme04320) and the mTOR signaling pathway (KEGG ID: dme04150).

## Discussion

The present study used a *Drosophila* model to investigate the extent to which anatomical sexual dimorphism impacts sex differences in the metabolome. We measured 91 targeted metabolomic features across three tissue types, comparing wildtype with *dsx* null flies, which exhibit reduced sexual dimorphism. Our findings underscore the importance of considering genetic mechanisms underlying sexual dimorphism when analyzing metabolic profiles.

The significant reduction in sex differences in metabolite levels observed in *dsx* null flies is a key finding of this study. Approximately half of the metabolites we measured showed significant sex differences in wildtype flies, while only one metabolite differed between *dsx* null sexes. The *dsx* gene plays a pivotal role in tipping the balance of gonad stem cells toward a “female” or “male” program during development (37), as well as directing sex-specific gene expression throughout adulthood that influences mating behaviors in both sexes (38–40). In the absence of functional *dsx*, the gonad and genital discs of *Drosophila* develop a morphology that is intermediate to typically developing XX and XY flies (26). We also observed a convergence of metabolic profiles between *dsx* null sexes, demonstrating how anatomical dimorphism can significantly influence the metabolome.

One notable exception to the trend of reduced sex differences in *dsx* null flies was kynurenate, a metabolite that consistently displayed higher levels in XX flies, regardless of the presence or absence of *dsx*, suggesting that sex-specific patterns of kynurenate are regulated by mechanisms independent of *dsx*. As *dsx* is an autosomal gene, its influence on sex-specific metabolism can be decoupled from the action of sex chromosomes. In our study, sex chromosome karyotypes were isogenic between SD groups, suggesting that the conserved sex differences in kynurenate are likely the result of an X- or Y-linked genetic factor. Two key genes at the top of the *Drosophila* tryptophan-kynurenine (Tryp-Kyn) degradation pathway are *vermilion* and *white* (41–43). Both genes are on the X chromosome (44,45), which may indicate a higher likelihood for X-linked sex biases in the metabolism of tryptophan and its derivatives, such as kynurenate. More research is needed to establish definitively whether the source of the sex difference in kynurenate here is X- or Y-linked.

The tissue-specificity of wildtype sex differences was another critical finding of this study. Different tissues exhibited unique sets of SD metabolites that were mostly eliminated in *dsx* null sexes. Clough *et al*. (27) identified that DSX protein isoforms bind thousands of the same targets across multiple tissues but result in sex- and tissue-specific functions. Rice *et al*. (28) later demonstrated that the expression of *dsx* in *Drosophila* is controlled by separate modular enhancers responsible for sex-specific traits in different organs. The variability in sex effects on metabolite levels across tissues thus could reflect the complexity with which DSX isoforms interact with diverse tissues, possibly to accommodate or compensate for sex-specific reproductive demands. Further work is needed to determine whether the influence of *dsx* on metabolite levels is a holdover from prior expression earlier in development to shape morphology, or if it has a more direct role in fine-tuning metabolic regulation.

We observed that the thorax tissue, which houses much of the fat body, shared more SD metabolites with both head and abdomen than head and abdomen shared with each other. This could reflect the fat body’s role as a key organ for inter-organ communication, regulating metabolism and developmental processes by releasing adipokines in response to nutritional and hormonal signals (46). *Transformer* (*tra*), a sex development gene that regulates the sex-specific splicing of *dsx* pre-mRNA (47), also regulates sex-specific body fat levels (4,5). It is unclear whether *dsx* plays a specific role downstream of *tra* in fat regulation or what specific organ it occurs in. However, Clough *et al*. (27) identified 25 genes in the fat body that were differently expressed in response to a switch in DSX isoform state between DSX^F^ and DSX^M^, suggesting that *dsx* is very active in the fat body. Lazareva *et al*. (48) hypothesized that the fat body is the source of secreted circulating proteins that reach the brain via hemolymph to drive male-typical courtship behaviors, similar to *dsx*-regulated mechanisms underlying Yolk Protein (49) and Collagen IV (50) secretions. Further experiments are needed to distinguish the roles of *tra*, *dsx* and downstream genes on metabolites in the fat body and other organs. As sex differences in the metabolome were reduced but not completely depleted in *dsx* null flies, some of the more minor sex effects could be due to *tra*.

Pathway enrichment for all SD metabolites revealed a significant enrichment of BCAA degradation and mTOR pathways. Sex differences in all three BCAAs (leucine, isoleucine and valine) were *dsx*-dependent in head and thorax tissues. BCAAs are potent activators of mTOR signaling (51), which is crucial for regulating various metabolic processes. The significant reduction in BCAA-related metabolites in our *dsx* null flies suggests that *dsx* plays a role in maintaining sex differences in BCAAs and, possibly, mTOR pathway activation as a result. This reduction in BCAA sex differences in the absence of *dsx* indicates that sex-specific regulation of the mTOR pathway is partly mediated by *dsx* or one of its downstream targets. The mTOR pathway also mediates the longevity effects of dietary restriction (52), which has sex-specific outcomes in *Drosophila* (53). In mice, restricting dietary BCAAs increased lifespan and metabolic health in males but not females (54). This supports the idea that sex-specific regulation of nutrient metabolism, including BCAAs, contributes to the differential effects of dietary restriction observed between sexes. Taking this one step further, our work supports the idea that anatomical dimorphism could influence these differential effects. Further research on the role of *dsx* in regulating BCAAs may provide insights into sex-related mechanisms that modify the response to *Drosophila* longevity interventions.

## Limitations

This study has several limitations that should be considered. These experiments were designed to distinguish the effect of sex, defined by karyotype, in a mutant where the external morphology of the sexes is nearly indistinguishable (26). To enable us to distinguish XX and XY flies without any clear external sexual dimorphism, we used a genetically marked Y-chromosome, Y-Bar[S], throughout the study. The bar-eyes phenotype manifests in eye morphology, which leads to a confound of eye morphology and karyotype. Thus, we acknowledge that some of the effects of sex on the metabolome, particularly in the head, may be due to eye morphology. If this eye morphology were to explain the sex difference in the metabolome, there is no known way in which these effects would depend on the *dsx* mutation. Similarly, as is common in *Drosophila* studies involving P-element-induced alleles, like the *dsx* null, the presence of the mini-*white* gene in the P-element, causes a confound between the *dsx* genotype and the presence of mini-white. We note that *white* is a component of the Tryp-Kyn pathway and so, expression of mini-*white* may influence metabolism. However, of all metabolites, sex differences in kynurenate were consistent across all tissues regardless of the presence of the P- element. Last, the study examined head, thorax, and abdomen tissues, each of which includes multiple organs. Our tissue-specific results cannot be attributed to any single organ.

## Conclusion

The findings from this study have several important implications for our understanding of variation in and regulation of physiological sex differences. First, they highlight the necessity of accounting for genetic mechanisms underlying sexual dimorphism, such as the role of the *doublesex* gene in *Drosophila*. The absence of *dsx* accompanies a significant reduction in sex differences in the metabolome, indicating that genetic context is crucial for understanding and interpreting sex differences in metabolic profiles. Second, the tissue-specific nature of *dsx* influence on the metabolome suggests that different tissues have unique metabolic demands and are differentially influenced by sex development gene networks. Comprehensive tissue-specific analyses can reveal insights into how sex differences manifest in various organs and how each organ might influence sex differences systemically. In conclusion, this study highlights the critical need for metabolomics research to incorporate genetic and phenotypic diversity related to sex characteristics.

## Methods

### Fly stocks and husbandry

We used two stocks containing *dsx* null deletions. Stock 1, *dsx*^f00683-d07058^ was obtained from the Bloomington *Drosophila* Stock Center with full genotype: *w*\*; P+PBac{*w*[+mC]=XP3.WH3}*dsx*^f000683-d07058^/*TM6B*, *Tb*^1^ [stock #66710]. To achieve isogenic lines, we first backcrossed this stock to our lab *w*^1118^ strain for five generations, tracking the *dsx* allele using the mini-*white* eye marker. As we did not observe viable homozygous *dsx* null offspring in this stock after crossing this line back to itself, we utilized a trans-heterozygous crossing scheme between our *dsx*^f000683-d07058^ strain and Stock 2, *dsx* deletion stock gifted from the lab of Mark Siegal. This second stock, *dsx*^f01649-d09625^, had been previously crossed in the Siegal lab to a dominant-Bar[S]-marked Y chromosome in a *w\** background and carried an *ix* mutation balanced by *CyO* with full genotype: *w*\**/*B[S]Y*; ix*[GFP]/*CyO*; *Df(3R)f01649-d09625/TM6B, Tb*.

We crossed Stock 2 to our previously backcrossed Stock 1 and observed the expected proportions of *dsx* null flies relative to wildtype flies. For our metabolomics experiment, we discarded flies with straight wings to remove the *ix* mutation from our experimental samples, and discarded flies with *TM6* to ensure that wildtype flies had an isogenic third chromosome. Thus, among the progeny of this cross, we recovered the four experimental genotypes (Table 4).

**Table 4.** Experimental Genotypes used in this study.

| Group | Eye Marker | Genotype |  |  |
| --- | --- | --- | --- | --- |
|  |  | Chrom 1 | Chrom 2 | Chrom 3 |
| XX – wildtype | *White eyes* | $\frac{w^{1118}}{w^*}$ | $\frac{+}{CyO}$ | $\frac{dsx^{f01649-d09625}}{+}$ |
| XY – wildtype | *White Bar eyes* | $\frac{w^{1118}}{B[S]Y}$ | $\frac{+}{CyO}$ | $\frac{dsx^{f01649-d09625}}{+}$ |
| XX – *dsx* null | *Orange eyes* | $\frac{w^{1118}}{w^*}$ | $\frac{+}{CyO}$ | $\frac{dsx^{f01649-d09625}}{dsx^{f00683-d07058}}$ |
| XY – *dsx* null | *Orange Bar eyes* | $\frac{w^{1118}}{B[S]Y}$ | $\frac{+}{CyO}$ | $\frac{dsx^{f01649-d09625}}{dsx^{f00683-d07058}}$ |

### Fly media and culture conditions

Flies were raised on a banana-based medium as described in (35) and housed in an incubator on a 12-hour light-dark cycle at 25°C.

### Tissue collection and metabolite extraction

All flies for metabolomics were collected three days post-eclosion and sorted with a two-minute timed exposure to CO2 into the four genotypes based on the segregating markers: orange eyes (mini-white), curly wings, and bar eyes. Flies were then allowed to recover for 24 hours in vials on fresh media before being flash-frozen in liquid nitrogen and stored at -80°C.

Individual flies were sectioned in a chilled petri dish on a cold metal block in dry ice, into head, thorax and abdomen samples using a chilled clean razor blade and sorted into 1.5mL Eppendorf tubes for storage at -80°C. Each sample consisted of tissue from 10 flies, with four to five biological replicates per tissue per genotype.

For metabolite extraction, one 5mm zirconium oxide bead was added to each Eppendorf tube before placing them in a frozen homogenizer block (Tissuelyser II, Qiagen). The tissue was pulverized by shaking at 30Hz for two minutes in a cold room. Samples were then suspended in 1 mL of a methanol:H2O 4:1 solution kept on dry ice, after which each sample was vortexed for 10 seconds. Tubes were then centrifuged at 14,000 rpm for 15 minutes at 4°C. 600 µL of supernatant was transferred to a new 1.5mL Eppendorf tube, dried under vacuum at 30°C overnight, and stored at -80°C when retrieved the following morning.

### Liquid chromatography-mass spectrometry (LC- MS)

Targeted LC-MS was carried out as described previously (18), providing measures of 91 metabolites.

### Statistical Methods

All statistical analysis was performed using R (version 4.3.0) open-source statistical software. Metabolite data were stratified by tissue (head, thorax, abdomen), then log-transformed, centered and scaled to have a mean = 0 and σ = 1 by sample. Principal component analysis (PCA) was then performed using the ‘prcomp’ function in R.

## Outlier Detection

One replicate sample of XX-wildtype/abdomen (sample XXwA4) was reported by LC-MS Core staff as having aberrant values for several compounds and thus investigated as a potential outlier using the ‘boxplot.stats’ function in R. This function identifies outliers as values that fall beyond 1.5 times the interquartile range (IQR) from the first and third quartiles. Sample XXwA4 had 8 outlier metabolite values by this method, the most of all samples. We ran analyses both with and without this sample and found that when the sample was excluded, more metabolites were significantly different in abdomen between wildtype sexes at FDR<5% (30 vs 24). The complete 51 metabolome samples are included in our main analysis (17 head, 17 thorax and 17 abdomen). Results of the abdomen analysis excluding sample XXwA4 are reported in Supplementary File 2.

### Univariate Analysis

To identify metabolites whose effect sizes differed between the four experimental genotypes (EG) in Table 4, we first performed one-way ANOVA on each normalized metabolite within each tissue type (head, thorax, abdomen), using the ‘lm’ function in R, followed by post-hoc analysis using the ‘TukeyHSD’ function in R to retrieve effect sizes between XX and XY samples for each SD group. P-values from the TukeyHSD were adjusted for multiple comparisons within each tissue using the ‘p.adjust’ function in R with method = “fdr”. Metabolites that met an FDR<5% were considered significantly different between the sexes in either wildtype, *dsx* null or both.

To determine whether there was a significant reduction in sex effect sizes across metabolites in each SD group, effect sizes between XX and XY from the Tukey HSD were compared between wildtype and *dsx* null by modeling the log-transformed absolute effect size as a function of the SD Group. P-values for these comparisons were derived from the ‘anova’ function in R on the linear models for each tissue.

### Metabolite Enrichment

Pathway analysis was conducted using the ‘FELLA’ package in R (36), which utilizes a network diffusion method to detect nodes in a biological network that are enriched for connectivity to small groups of metabolites, such as the 91 targeted metabolites measured in this study. Within FELLA, we accessed the *Drosophila melanogaster* KEGG Database (Release 111.0+). KEGG IDs were available for 85 of the 91 metabolites measured here. Enrichment of nodes within the KEGG network by a sub-set of the metabolites measured here, such as the metabolites with significant sex effects in our data, was tested by permuting among the 85 metabolites 10,000 times to give empirical P values. We corrected for multiple testing by applying the ‘p.adjust’ function in R with method = “fdr”.

## Acknowledgments

We kindly thank Mark Siegal and Carmen Robinett for the gift and permission to use the *dsx*^f01649-d09625^ fly strain, and Michelle Arbeitman and Barbara Wakimoto for additional advice on identifying *dsx* null phenotypes. This study used stock 66710 from the Bloomington *Drosophila* Stock Center (NIH P40OD018537). Many additional thanks to Elizabeth Rideout and Harmit Malik for reading and commenting on this manuscript in early stages. RC was supported by the Mary Gates Endowment and NIH T32 AG066574. DP received support from USDA cooperative agreement USDA/ARS 58-8050-9-004. This project was also supported by NIA R21 AG074495 and NIH P30 AG013280.

## Data Accessibility

Raw metabolome data and the R code used for analyses and figure generation for this manuscript are available for download on GitHub: https://github.com/rcoig/dsx-Metabolomics.

## Abbreviations

dsx: doublesex
SD: sex difference
tra: transformer
Tryp-Kyn: tryptophan-kynurenine
w: white

## References

1. Ober C, Loisel DA, Gilad Y. Sex-specific genetic architecture of human disease. Nat Rev Genet. 2008;9(12):911–22.

2. Mauvais-Jarvis F, Merz NB, Barnes PJ, Brinton RD, Carrero JJ, DeMeo DL, et al. Sex and gender: modifiers of health, disease, and medicine. Lancet Lond Engl. 2020;396(10250):565–82.

3. Klein SL, Flanagan KL. Sex differences in immune responses. Nat Rev Immunol. 2016;16(10):626–38.

4. Wat LW, Chowdhury ZS, Millington JW, Biswas P, Rideout EJ. Sex determination gene transformer regulates the male-female difference in *Drosophila* fat storage via the adipokinetic hormone pathway. Elife. 2021;10:e72350.

5. Wat LW, Chao C, Bartlett R, Buchanan JL, Millington JW, Chih HJ, et al. A role for triglyceride lipase brummer in the regulation of sex differences in *Drosophila* fat storage and breakdown. Plos Biol. 2020;18(1):e3000595.

6. Mank JE, Rideout EJ. Developmental mechanisms of sex differences: from cells to organisms. Development. 2021;148(19).

7. Millington JW, Brownrigg GP, Chao C, Sun Z, Basner-Collins PJ, Wat LW, et al. Female-biased upregulation of insulin pathway activity mediates the sex difference in *Drosophila* body size plasticity. Elife. 2021;10:e58341.

8. Millington JW, Rideout EJ. Sex differences in *Drosophila* development and physiology. Curr Opin Physiology. 2018;6:46–56.

9. McCarthy MM, Arnold AP, Ball GF, Blaustein JD, Vries GeertJD. Sex Differences in the Brain: The Not So Inconvenient Truth. J Neurosci. 2012;32(7):2241–7.

10. Johnson CH, Ivanisevic J, Siuzdak G. Metabolomics: beyond biomarkers and towards mechanisms. Nat Rev Mol Cell Biol. 2016;17(7):451–9.

11. Zampieri M, Sauer U. Metabolomics-driven understanding of genotype-phenotype relations in model organisms. Curr Opin Syst Biol. 2017;6:28–36.

12. Rohde PD, Kristensen TN, Sarup P, Muñoz J, Malmendal A. Prediction of complex phenotypes using the *Drosophila* melanogaster metabolome. Heredity. 2021;126(5):717–32.

13. Hoffman JM, Soltow QA, Li S, Sidik A, Jones DP, Promislow DEL. Effects of age, sex, and genotype on high-sensitivity metabolomic profiles in the fruit fly, *Drosophila* melanogaster. Aging Cell. 2014;13(4):596–604.

14. Zhou S, Morgante F, Geisz M, Ma J, Anholt R, Mackay T. Systems Genetics of the *Drosophila* Metabolome. Genome Res. 2019;30(3):gr.243030.118.

15. Chintapalli VR, Bratty MA, Korzekwa D, Watson DG, Dow JAT. Mapping an Atlas of Tissue-Specific *Drosophila* melanogaster Metabolomes by High Resolution Mass Spectrometry. Plos One. 2013;8(10):e78066.

16. Harrison BR, Lee MB, Zhang S, Young B, Han K, Sukomol J, et al. Wide-ranging genetic variation in sensitivity to rapamycin in *Drosophila* melanogaster. Aging Cell. 2024;e14292.

17. Jin K, Wilson KA, Beck JN, Nelson CS, Brownridge GW, Harrison BR, et al. Genetic and metabolomic architecture of variation in diet restriction-mediated lifespan extension in *Drosophila*. Plos Genet. 2020;16(7):e1008835.

18. Zhao X, Golic FT, Harrison BR, Manoj M, Hoffman EV, Simon N, et al. The metabolome as a biomarker of aging in *Drosophila* melanogaster. Aging Cell. 2022;21(2):e13548.

19. Harrison BR, Wang L, Gajda E, Hoffman EV, Chung BY, Pletcher SD, et al. The metabolome as a link in the genotype-phenotype map for peroxide resistance in the fruit fly, *Drosophila* melanogaster. Bmc Genomics. 2020;21(1):341.

20. Soller M, Bownes M, Kubli E. Control of Oocyte Maturation in Sexually Mature*Drosophila*Females. Dev Biol. 1999;208(2):337–51.

21. Salz H, Erickson JW. Sex determination in *Drosophila*: The view from the top. Fly. 2010;4(1):60–70.

22. Saccone G. A history of the genetic and molecular identification of genes and their functions controlling insect sex determination. Insect Biochem Mol Biol. 2022;151:103873.

23. Baker BS, Wolfner MF. A molecular analysis of doublesex, a bifunctional gene that controls both male and female sexual differentiation in *Drosophila* melanogaster. Genes Dev. 1988;2(4):477–89.

24. Burtis KC, Baker BS. *Drosophila* doublesex gene controls somatic sexual differentiation by producing alternatively spliced mRNAs encoding related sex-specific polypeptides. Cell. 1989;56(6):997–1010.

25. Coschigano KT, Wensink PC. Sex-specific transcriptional regulation by the male and female doublesex proteins of *Drosophila*. Genes Dev. 1993;7(1):42–54.

26. Hildreth PE. Doublesex, a recessive gene that transforms both males and females of *Drosophila* into intersexes. Genetics. 1965;51(4):659–78.

27. Clough E, Jimenez E, Kim YA, Whitworth C, Neville MC, Hempel LU, et al. Sex- and Tissue-Specific Functions of *Drosophila* Doublesex Transcription Factor Target Genes. Dev Cell. 2014;31(6):761–73.

28. Rice GR, Barmina O, Luecke D, Hu K, Arbeitman M, Kopp A. Modular tissue-specific regulation of doublesex underpins sexually dimorphic development in *Drosophila*. Development. 2019;146(14):dev178285.

29. Robinett CC, Vaughan AG, Knapp JM, Baker BS. Sex and the Single Cell. II. There Is a Time and Place for Sex. Plos Biol. 2010;8(5):e1000365.

30. Barmina O, Gonzalo M, McIntyre LM, Kopp A. Sex- and segment-specific modulation of gene expression profiles in *Drosophila*. Dev Biol. 2005;288(2):528–44.

31. Telonis-Scott M, Kopp A, Wayne ML, Nuzhdin SV, McIntyre LM. Sex-Specific Splicing in *Drosophila*: Widespread Occurrence, Tissue Specificity and Evolutionary Conservation. Genetics. 2009;181(2):421–34.

32. Arbeitman MN, New FN, Fear JM, Howard TS, Dalton JE, Graze RM. Sex Differences in *Drosophila* Somatic Gene Expression: Variation and Regulation by doublesex. G3 Genes Genomes Genetics. 2016;6(7):1799–808.

33. Lee G, Hall JC, Park JH. Doublesex gene expression in the central nervous system of *Drosophila melanogaster*. J Neurogenet. 2002;16(4):229–48.

34. Rideout EJ, Dornan AJ, Neville MC, Eadie S, Goodwin SF. Control of sexual differentiation and behavior by the doublesex gene in *Drosophila* melanogaster. Nat Neurosci. 2010;13(4):458–66.

35. Harrison BR, Hoffman JM, Samuelson A, Raftery D, Promislow DEL. Modular Evolution of the *Drosophila* Metabolome. Mol Biol Evol. 2021;39(1):msab307.

36. Picart-Armada S, Fernández-Albert F, Vinaixa M, Yanes O, Perera-Lluna A. FELLA: an R package to enrich metabolomics data. Bmc Bioinformatics. 2018;19(1):538.

37. Camara N, Whitworth C, Dove A, Doren MV. Doublesex controls specification and maintenance of the gonad stem cell niches in *Drosophila*. Development. 2019;146(11):dev170001.

38. Pan Y, Robinett CC, Baker BS. Turning Males On: Activation of Male Courtship Behavior in *Drosophila* melanogaster. PLoS ONE. 2011;6(6):e21144.

39. Rezával C, Nojima T, Neville MC, Lin AC, Goodwin SF. Sexually Dimorphic Octopaminergic Neurons Modulate Female Postmating Behaviors in *Drosophila*. Curr Biol. 2014;24(7):725–30.

40. Rezával C, Pattnaik S, Pavlou HJ, Nojima T, Brüggemeier B, D’Souza LAD, et al. Activation of Latent Courtship Circuitry in the Brain of *Drosophila* Females Induces Male-like Behaviors. Curr Biol. 2016;26(18):2508–15.

41. Green MM. A study of tryptophane in eye color mutants of *Drosophila*. Genetics. 1949;34(5):564–72.

42. Rizki TM. Mutant genes regulating the indicibility of kynurenine synthesis. J Cell Biol. 1964;21(2):203–11.

43. Mackenzie SM, Brooker MR, Gill TR, Cox GB, Howells AJ, Ewart GD. Mutations in the white gene of *Drosophila* melanogaster affecting ABC transporters that determine eye colouration. Biochim Biophys Acta (BBA) - Biomembr. 1999;1419(2):173–85.

44. Walker AR, Howells AJ, Tearle RG. Cloning and characterization of the vermilion gene of *Drosophila* melanogaster. Mol Gen Genet MGG. 1986;202(1):102–7.

45. Levis R, Bingham PM, Rubin GM. Physical map of the white locus of *Drosophila* melanogaster. Proc Natl Acad Sci. 1982;79(2):564–8.

46. Meschi E, Delanoue R. Adipokine and fat body in flies: Connecting organs. Mol Cell Endocrinol. 2021;533:111339.

47. Hoshijima K, Inoue K, Higuchi I, Sakamoto H, Shimura Y. Control of doublesex Alternative Splicing by transformer and transformer-2 in *Drosophila*. Science. 1991;252(5007):833–6.

48. Lazareva AA, Roman G, Mattox W, Hardin PE, Dauwalder B. A Role for the Adult Fat Body in *Drosophila* Male Courtship Behavior. Plos Genet. 2007;3(1):e16.

49. Belote JM, Handler AM, Wolfner MF, Livak KJ, Baker BS. Sex-specific regulation of yolk protein gene expression in *Drosophila*. Cell. 1985;40(2):339–48.

50. Peng Q, Chen J, Wang R, Zhu H, Han C, Ji X, et al. The sex determination gene doublesex regulates expression and secretion of the basement membrane protein Collagen IV. J Genet Genom. 2022;49(7):636–44.

51. Juricic P, Grönke S, Partridge L. Branched-chain amino acids have equivalent effects to other essential amino acids on lifespan and ageing-related traits in *Drosophila*. Journals Gerontology Ser. 2019;75(1):glz080-.

52. Katewa SD, Kapahi P. Role of TOR signaling in aging and related biological processes in *Drosophila* melanogaster. Exp Gerontol. 2011;46(5):382–90.

53. Magwere T, Chapman T, Partridge L. Sex Differences in the Effect of Dietary Restriction on Life Span and Mortality Rates in Female and Male *Drosophila* Melanogaster. Journals Gerontology Ser. 2004;59(1):B3–9.

54. Richardson NE, Konon EN, Schuster HS, Mitchell AT, Boyle C, Rodgers AC, et al. Lifelong restriction of dietary branched-chain amino acids has sex-specific benefits for frailty and lifespan in mice. Nat Aging. 2021;1(1):73–86.

