## Supplementary figures and images for "Tissue-specific metabolomic signatures for a *doublesex* model of reduced sexual dimorphism"

### Supplementary Fig 1

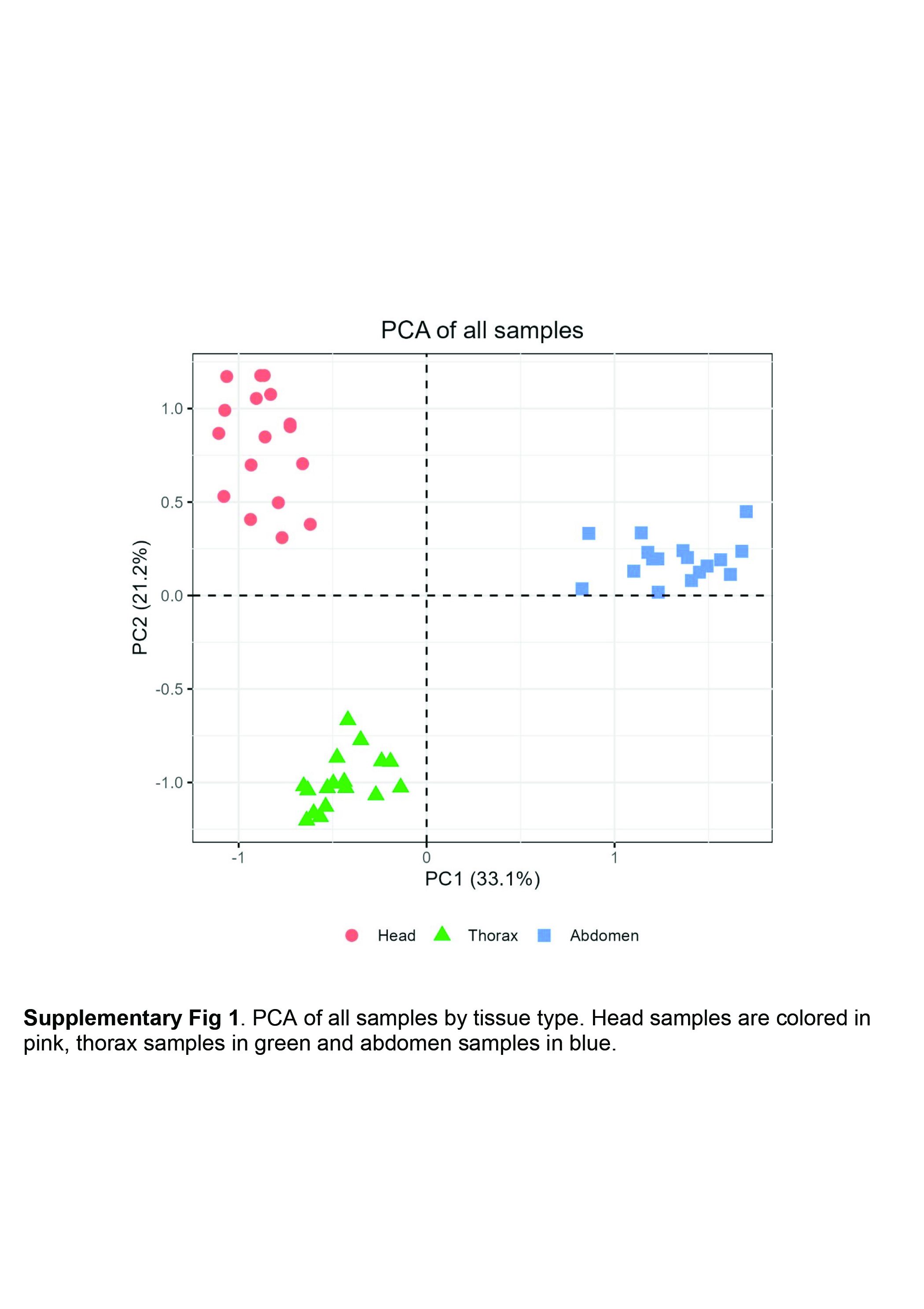
